# *Giardia* Increases Macrophage Production of the Anti-Inflammatory Cytokine Interleukin-10 in Response to Lipopolysaccharide via Macrophage Galactose Binding Lectin (MGL1)

**DOI:** 10.1101/2024.04.15.589568

**Authors:** V Angelova, M Darmadi, E Miskovsky, M Fink, S Menegas, H Wexelblatt, S Singer

**Affiliations:** Georgetown University

## Abstract

*Giardia duodenalis* is an intestinal protozoan parasite, common in low- and middle-income countries. Infection is frequently subclinical, even when it is associated with other pathologies like growth stunting in children. Recent longitudinal cohort studies have found *Giardia* more frequently in patients with milder symptoms and have even suggested that *Giardia* reduces rotavirus symptom severity. One potential mechanism for limiting disease severity due to other enteropathogens is the promotion of anti-inflammatory responses that limit pathology. Our lab previously showed that *Giardia* reduced production of IL-12 by dendritic cells stimulated with TLR agonists. In this study, we show that *Giardia* increases the production of the anti-inflammatory cytokine IL-10 by mouse peritoneal macrophages in response to bacterial LPS. This potentiation is specific to IL-10, as no changes were seen in the production of the pro-inflammatory cytokine TNF-□. Moreover, peritoneal macrophages from mice lacking macrophage galactose-binding lectin (MGL1), a pathogen recognition receptor that has been previously shown to bind N-acetylgalactosamine, failed to increase IL-10 production after stimulation with *Giardia* and LPS. *Giardia’*s immunoregulation of the IL-10 response may help us understand the parasite’s role in reducing diarrheal severity.

## INTRODUCTION

*Giardia duodenalis* is an intestinal protozoan parasite, most commonly transmitted via contaminated water. There are an estimated 280 million cases annually worldwide, most of which occur in low- and middle-income countries. Infection may have a subclinical presentation or may cause common intestinal distress symptoms, including diarrhea, cramps, and bloating (Wolfe 1992). Even though giardiasis in healthy individuals resolves on its own or with metronidazole treatment, post-infectious sequelae are observed in many patients (Halliez and Buret 2013). The Malnutrition and Enteric Disease study (MAL-ED) identified that *Giardia* is one of the five leading causes of growth stunting in children under two (Rogawski et al. 2018) and has been epidemiologically linked to cognitive and motor developmental delays (Berkman et al. 2002; Budge et al. 2019). Importantly, stunting was seen regardless of the presence of giardiasis-induced diarrhea.

*Giardia* notably does not cause acute, life-threatening diarrhea in children; the mechanisms behind this observation warrant further study (Kotloff et al. 2013; Muhsen and Levine 2012; Platts-Mills et al. 2015). Meta-analysis of the association between giardiasis and acute diarrhea (3 loose stools in under 24 hours) identified a positive link in infants younger than 3 months of age but not in any other age group. In several studies, in fact, individuals infected with *Giardia* had a lower risk of acute diarrhea than controls (Muhsen and Levine 2012). The extensive Global Enteric Multicenter Study (GEMS) similarly showed that *Giardia* was not a contributor to moderate-to-severe diarrhea and was more commonly found in control stools than in patient stools (Kotloff et al. 2013).

In the small intestine, *Giardia* interacts with many bacteria, both commensal microbiome members and other pathogenic strains (Fink and Singer 2017). *Giardia* has been shown to interact with bacterial lipopolysaccharide (LPS) to limit production of the pro-inflammatory cytokine IL-12 and promote the secretion of the anti-inflammatory cytokine IL-10 from co-stimulated bone marrow-derived dendritic cells (BMDC) (Kamda and Singer 2009). Further, BMDCs from *Giardia*-infected mice fed a low protein diet have an increased production of IL-10 in response to LPS compared to BMDCs from uninfected animals or infected animals fed a regular diet (Burgess et al. 2019). While the primary role of dendritic cells is to present antigens to T cells, peritoneal macrophages are effectors tasked with intestinal surveillance (Yip et al. 2021). There are currently no studies on the macrophage response to dual stimulations with *Giardia* and LPS, even though these innate cells proliferate significantly in the lamina propria post infection (Fink et al. 2019).

There are similarly very few studies identifying the host receptors through which the immune system can detect the presence of *Giardia* signals. It has previously been shown that *Giardia* binds to the mannose binding lectin (MBL) activating the complement pathway, with the interaction likely between MBL and N-acetylglucosamine (GlcNAc) on the surface of the *Giardia* trophozoite (Li, Tako, and Singer 2016). GlcNAc is a product of the hexosamine pathway and has been frequently reported as a glycosylation agent on *Giardia* trophozoites and excretory-secretory products (Banerjee, Robbins, and Samuelson 2009; Das and Gillin 1996; Morelle et al. 2005; Ratner et al. 2008; Ward et al. 1988). In contrast to trophozoites, the cyst wall of *Giardia* contains an abundant carbohydrate polymer composed of β_1-3_ linked N-acetylgalactosamine (GalNAC), which could potentially signal through macrophage galactose-type lectin (MGL) receptors (Gerwig et al. 2002). The potential interaction between *Giardia* and MGL has not been explored. Notably, humans have a single gene for the MGL receptor on macrophages and dendritic cells; while mice generally express MGL1 on macrophages, pDCs, and cDCs, MGL2 is mostly found on cDCs (Denda-Nagai et al. 2010).

In this study we focused on the role of *Giardia*-induced MGL1 signaling on murine peritoneal macrophages in potentiating the IL-10 response to bacterial LPS. An increase of anti-inflammatory cytokines in response to MGL1 and TLR4 co-stimulation could help explain the findings of reduced diarrheal burden yet the presence of *Giardia* in control stools (Kotloff et al. 2013). Notably, MGL1 has been previously shown to upregulate the IL-10 response in peritoneal macrophages (Kurashina et al. 2021). Additionally, MGL1 and TLR2 activation have been shown to synergize in the production of IL-10 and TNF-□ (van Vliet et al. 2013).

## MATERIALS AND METHODS

### Parasite Extracts

*Giardia duodenalis* clone H7 assemblage GS (ATCC# 50581) trophozoites were grown to confluence in TYI-S-33 media and detached on ice (Aggarwal and Nash 1987). Parasites collected from 200-250 ml of confluent cultures were centrifuged at 1000 RCF for 10 minutes, washed 3-4 times, and resuspended in 1 mL of sterile, cold PBS. Concentrated parasites were then lysed with 10-20 repetitive cycles of freeze-thawing. Extract protein was quantified by measuring absorbance at 280 nm using a nanodrop (Thermo Scientific, NanoDrop 2000c).

In some experiments the extract was heated at 65º for 30 minutes or centrifuged at 18,000 RCF for 5 minutes before being added to macrophage cultures.

### Animals

Adult C57BL/6J mice (wildtype) were purchased from Jackson laboratories. MGL1^-/-^ (B6.129-*Clec10a*^*tm1Hed*^/J; (Onami et al. 2002) breeders were provided by Dr. Jyotika Sharma from the University of North Dakota and bred at Georgetown University’s Division of Comparative Medicine. After crossing MGL1^-/-^ mice and C57BL/6J mice to produce heterozygous F1 mice, MGL1^+/-^ mice were backcrossed to MGL1^-/-^ to produce homozygous knockout mice and heterozygous littermate controls for experiments J. All animals were maintained in specific pathogen free conditions.

### Isolation of peritoneal macrophages

Peritoneal macrophages (PM) were isolated from adult mice using previously published methods (Ray and Dittel 2010). Briefly, mice were euthanized using CO_2_ and cervical dislocation following IACUC protocols. The skin covering the abdominal cavity was retracted, and the peritoneum was exposed. Sterile PBS was injected into the peritoneal cavity. The peritoneum was massaged, and the PBS carrying peritoneal macrophages was collected using a syringe. Cells were centrifuged at 300 RCF for 5 minutes, resuspended in 1 mL of sterile macrophage media, and counted.

### Peritoneal macrophage culture

Peritoneal macrophages were plated at a concentration of 1x10^6^/mL in 96-well plates (individual animals for MGL1^+/-^ and MGL1^-/-^ experiments) or 24-well plates (pooled samples for all remaining experiments). Pools consisted of peritoneal macrophages from 2-3 animals. Cells were cultured in Dubecco’s Modified Eagle Medium (Thermo Fisher Scientific) supplemented with 10% fetal bovine serum (Omega Scientific, Inc.), 55 μM 2-mercaptoethanol (Gibco), and 1% antibiotic-antimycotic solution (Gibco). Macrophages were allowed to adhere overnight in a 37°C, 5% CO_2_ incubator before stimulation.

Media was removed and macrophages were stimulated for 24 hours with 10 ng/mL *Salmonella typhimurium* LPS (Invivogen), 100 μg/mL *Giardia* extract, or a combination of both in fresh media.

### Cytokine quantification in supernatants

Supernatants (1 mL from individual wells in 24-well plates and 250 μL from individual wells in 96-well plates) were collected and directly used for ELISAs (DuoSet® ELISA kits from R&D Systems). Supernatants from LPS conditions were diluted 1:5 for IL-10 quantification and 1:10 for TNF-□ quantification to avoid reaching the upper limit of detection. Measurements below the limit of detection were set to the limit for graphing and statistical analyses. Samples and standard curves were measured in duplicate.

### Calculation of IL-10 potentiation ratio

The 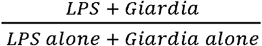 ratio for IL-10 was calculated for every pool of wildtype mice; this ratio is defined as the “IL-10 potentiation ratio” throughout the paper. The IL-10 potentiation ratio was then compared to expected additive ratio of 1 to determine if the potentiation is synergistic or additive.

### Data analysis

Data was plotted and analyzed using Prism. The appropriate statistical tests are indicated in the figure legends.

## RESULTS

### *Giardia* potentiates production of IL-10 but not TNF-□ in response to LPS stimulation

To characterize the peritoneal macrophage response to *Giardia*, we isolated macrophages from adult C57BL/6J mice and stimulated the cells *ex vivo*. Macrophages were stimulated with *Salmonella typhimurium* LPS, *Giardia* extract, or both LPS and *Giardia* extract (dual stimulation).

*Giardia* extract alone elicited almost no TNF-□ production from peritoneal macrophages; cytokine levels often fell below the limit of detection (Fig. 1a; Supplementary table 1). In contrast, macrophages stimulated with *Salmonella typhimurium* LPS or dual stimulated with LPS and *Giardia* extract produced very high, but indistinguishable amounts of TNF-L. The same trend was observed across our peritoneal macrophage pools (Fig. 1b). *Giardia* had no potentiating effect on the TNF-□ response to LPS.

**Figure 1:**
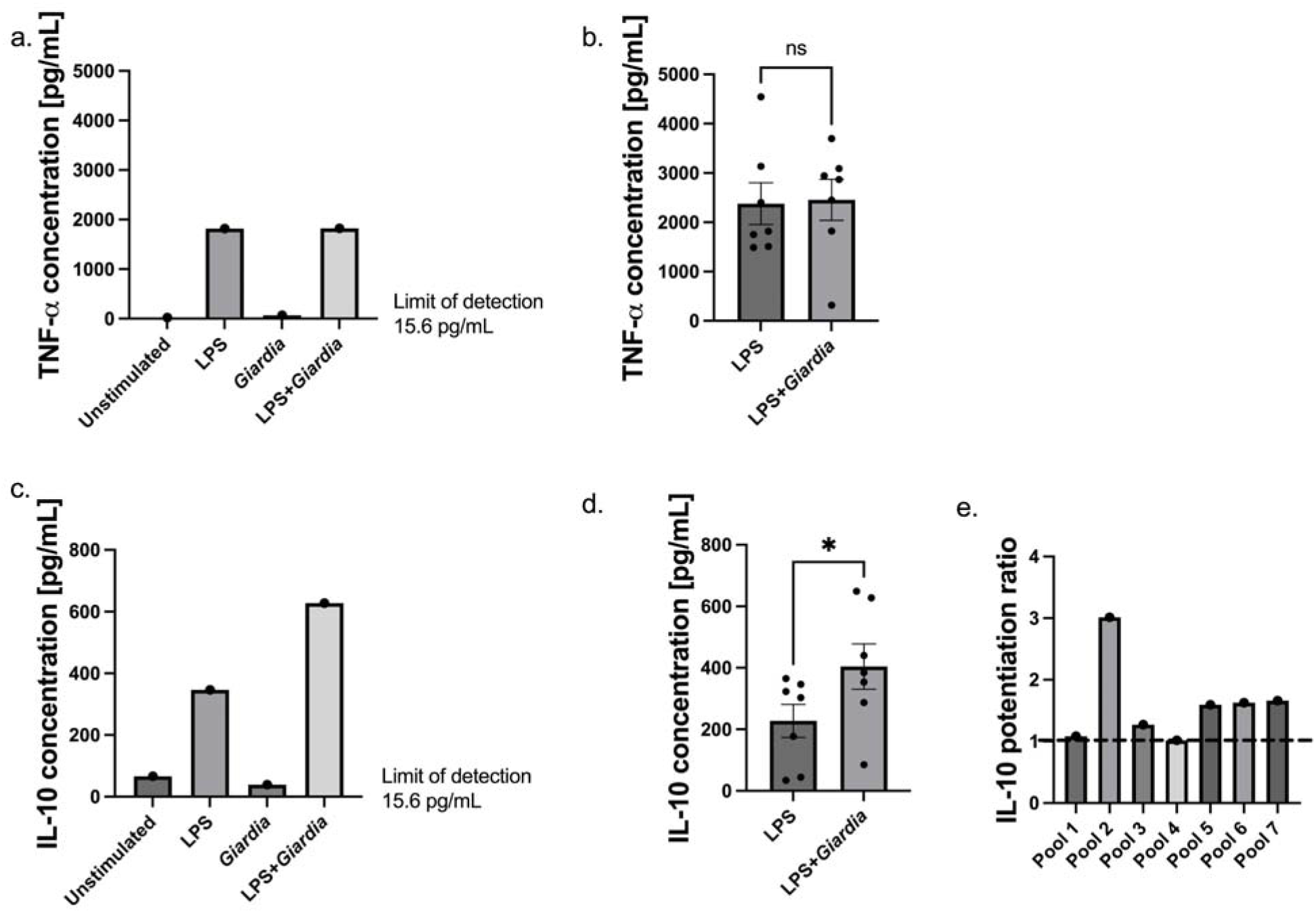
Dual stimulation of peritoneal macrophages with *Salmonella typhimurium* LPS and *Giardia duodenalis* extract potentiates IL-10 production. Peritoneal macrophages from C57BL/6J mice were plated and stimulated as described in the methods. Each data point is representative of 2-3 pooled animals. Macrophages were stimulated with 100 μg/mL *Giardia*, 10 ng/mL LPS, or both in fresh macrophage media 24 hours after plating. **a**. Representative TNF-□ ELISA data from one pool of peritoneal macrophages. **b**. Raw TNF-□ concentration values due to stimulation with LPS alone or dual stimulation. **c**. Representative IL-10 ELISA data from one pool of peritoneal macrophages. **d**. Raw IL-10 cytokine production from LPS alone or dual stimulation. **e**. Analysis of the IL-10 potentiation ratio. **b, d**. Data were analyzed with two-tailed Wilcoxon signed-rank test for paired, non-parametric datasets, *p<0.05.

*Giardia* extract alone similarly elicited little IL-10 production from the peritoneal macrophages (Fig. 1c; Supplementary Table 2). However, dual stimulation nearly doubled the IL-10 production compared to stimulation with LPS alone. Again, this trend was maintained across multiple pools, with an average IL-10 potentiation of about 2-fold (Fig. 1d). We were interested in whether this potentiation was due to additive or synergistic effects, so we calculated the IL-10 potentiation ratio for every pool of macrophages (Fig. 1e). The ratio was higher than the expected additive ratio of 1 in the majority of the pools, suggesting that *Giardia* and LPS were synergizing in the production of the anti-inflammatory IL-10. Note that the ratios for pool 1 and 4 are very close to 1 but over the cut-off.

### *Giardia* potentiation of IL-10 requires MGL1 signaling

Because MGL has been shown to enhance production of IL-10 from dendritic cells costimulated with the TLR1/2 ligand Pam3Csk4, we decided to test if this lectin was also involved in the enhanced production of IL-10 by macrophages seen when using Giardia extracts (van Vliet et al. 2013). We isolated peritoneal macrophages from MGL1^-/-^ and littermate control MGL1^+/-^ adult mice. Littermates were used to minimize the effects of the microbiome on the immune response. Cells from individual animals were plated and stimulated with LPS, Giardia extract, or both.

Macrophages from MGL1^+/-^ animals produced significantly more IL-10 when dual-stimulated with LPS and *Giardia* than with LPS alone (Figure 2a; Supplementary Table 3). Notably, the IL-10 potentiation in MGL1^+/-^ animals was lower than that seen in non-littermate control wildtype mice. The discrepancy could be due to a gene dosage effect, differences arising from testing pools of cells in 24-well plates instead of cells from individual mice in a 96-well format, or perhaps differences in the microbiome and breeding conditions between the wildtype animals that were purchased from Jackson Labs compared to the MGL1^+/-^ animals bred at Georgetown University. Importantly, macrophages from MGL1^-/-^ mice produced indistinguishable amounts of IL-10 following dual stimulation or stimulation with LPS alone, indicating that *Giardia* failed to potentiate the IL-10 response in MGL1^-/-^ macrophages (Fig. 2b; Supplementary table 4).

**Figure 2:**
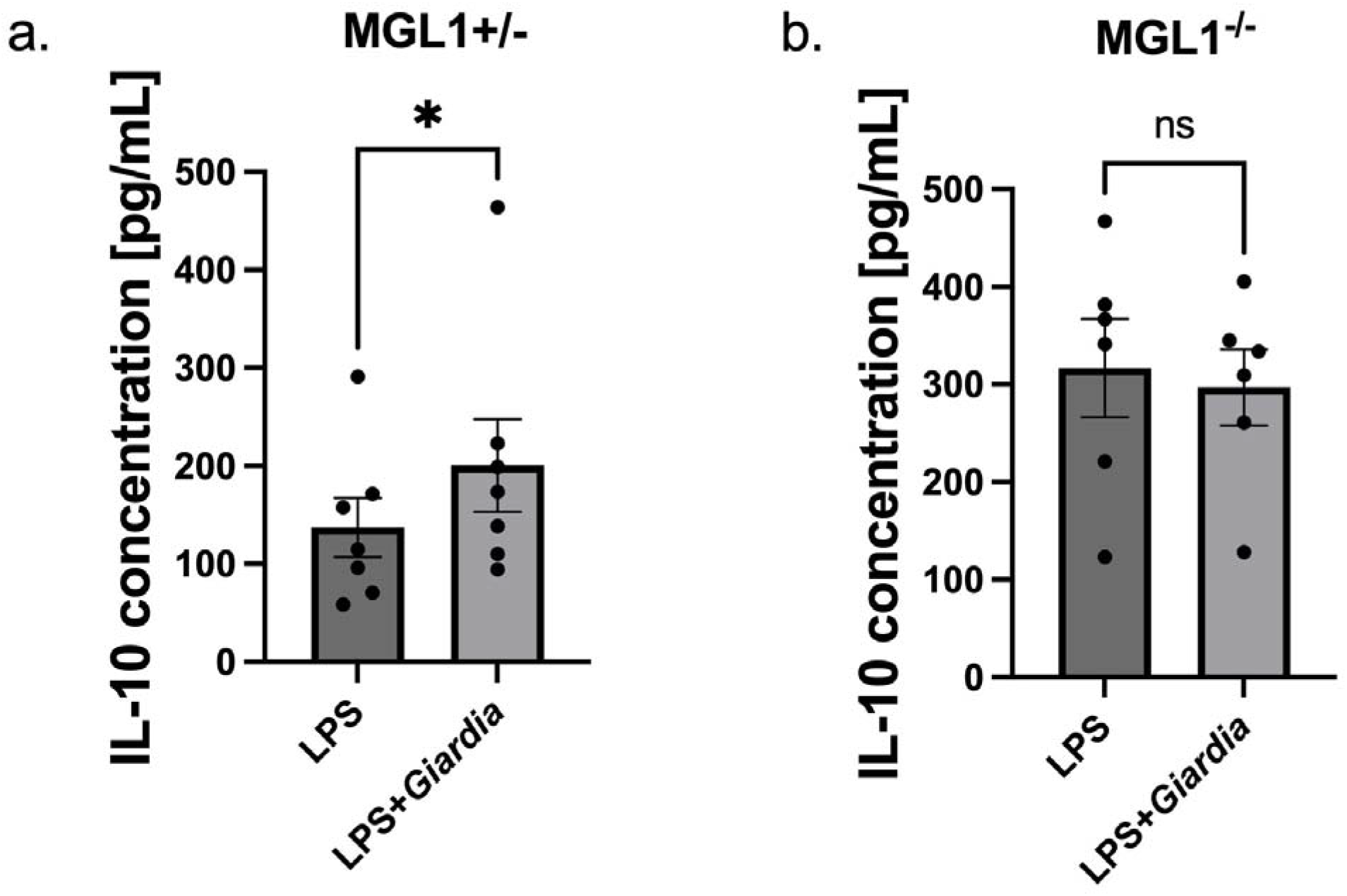
*Giardia*-induced IL-10 potentiation of the LPS response is dependent on the MGL1 receptor. Peritoneal macrophages were isolated from individual MGL1^+/-^ (**a**) and MGL1^-/-^ (**b**) mice were stimulated with 10 ng/mL LPS, 100 μg/mL *Giardia* extract, or both in fresh media 24 hours after plating. IL-10 production was measured via ELISA. Data were analyzed with two-tailed Wilcoxon signed-rank test for paired, non-parametric datasets, *p<0.05.

### Heating the *Giardia* extract affects its potentiation ability

To begin determining the nature of the MGL1 ligand on *Giardia*, we decided to either heat or centrifuge the parasite extract prior to stimulation. The goal of heating (65□ for 30 minutes) was to denature proteins in the extract, while centrifugation (18,000 RCF for 5 minutes) would help remove high molecular weight polymers. Previous research has identified that the *Giardia* cyst wall is composed of high molecular weight polymers of GalNAc, and this could be a potential ligand for MGL1 (Gerwig et al. 2002).

Peritoneal macrophages from C57BL/6J wildtype mice were stimulated with LPS, untreated *Giardia* extract, heated *Giardia* extract, centrifuged *Giardia* extract, or were dually stimulated with both LPS and each of the three extracts. There was no difference in IL-10 production between the three types of extracts when stimulating with extract alone (Fig. 3a; Supplementary table 2). Rather than decreasing IL-10 production, heating the *Giardia* extract (i.e., denaturing protein) significantly increased IL-10 production in dual stimulation conditions (Fig. 3b; Supplementary table 2). This suggests that the ligand for MGL1 does not require a specific protein conformation and that *Giardia* proteases could be degrading IL-10 in the untreated extract conditions. *Giardia* proteases have been reported to degrade IL-8 secreted from intestinal epithelial cells (Cotton, Bhargava, et al. 2014). In contrast to the effect of heat treatment, centrifugation of the extract had no effect on IL-10 production (Fig. 3b; Supplementary Table 2). While high-speed centrifugation could remove free long-chain polymers, it likely does not remove any *Giardia* proteins glycosylated with terminal Gal or GalNAc.

**Figure 3:**
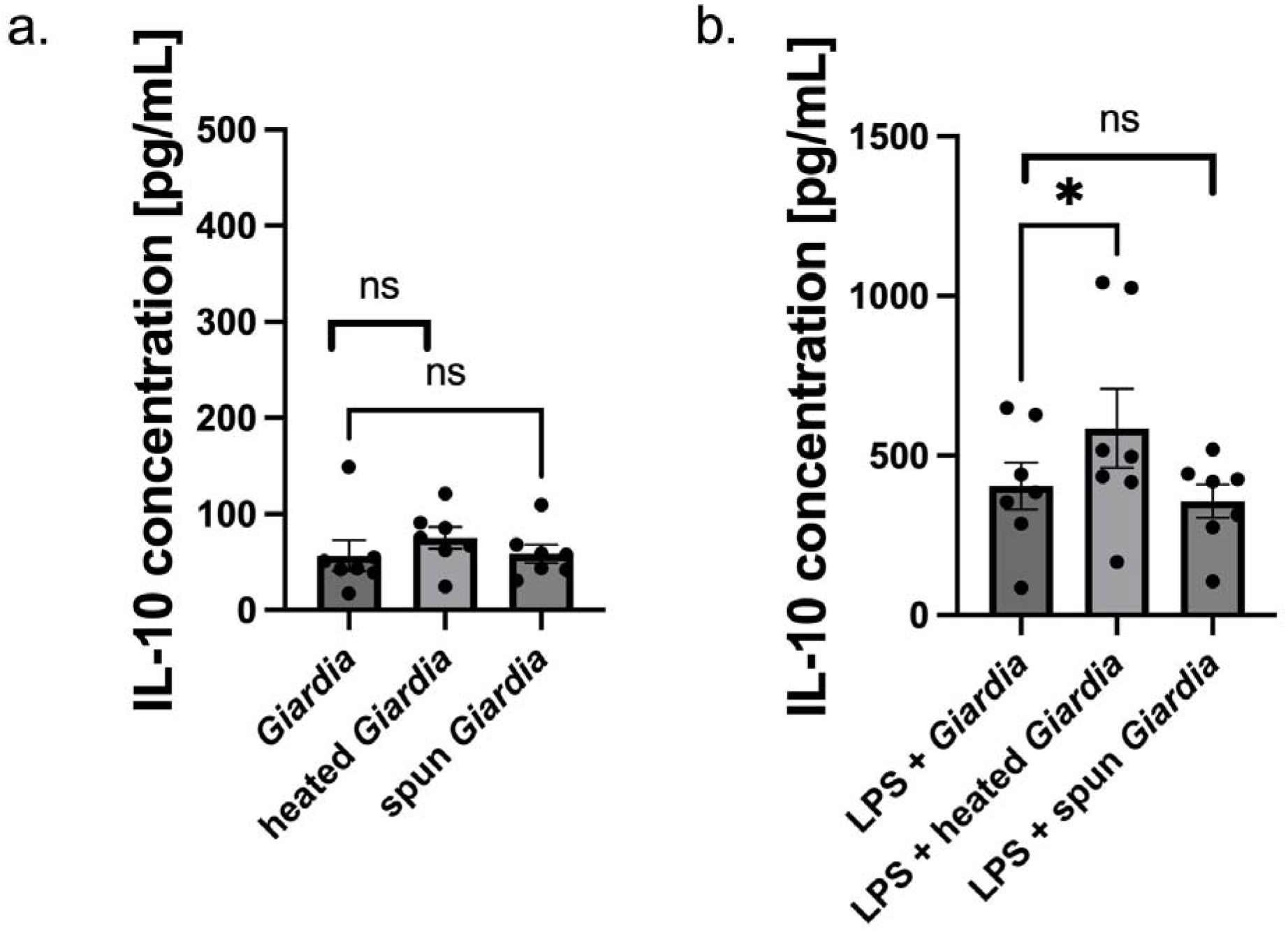
Heated *Giardia* extract amplifies IL-10 production in dual stimulation with LPS. Peritoneal macrophages from C57BL/6J adult mice were pooled and plated at 1x10^6^ cells/mL in 24-well plates. Cells were then stimulated with 10 ng/mL LPS and 100 μg/mL heated *Giardia* extract, centrifuged extract, or untreated extract. IL-10 levels in supernatants were quantified via ELISA. Note that these are the same pools of animals as the ones in Figure 1. **a**. IL-10 produced due to stimulation with each of the three *Giardia* extracts. **b**. IL-10 produced due to dual stimulation with LPS and each of the three *Giardia* extracts. Data were analyzed with two-tailed Mann-Whitney tests for paired, non-parametric data.

## DISCUSSION

In this study, we wanted to consider *Giardia*’s ability to modulate the macrophage immune response to bacterial LPS. We saw that *Giardia* and LPS synergize in the production of the anti-inflammatory cytokine IL-10 but not the pro-inflammatory cytokine TNF-alpha by peritoneal macrophages. The IL-10 potentiation depends on the MGL1 receptor, and is particularly interesting given *Giardia*’s failure to induce IL-10 on its own. Heating the *Giardia* extract prior to stimulation additionally enhanced the IL-10 production, likely due to the denaturation of parasite proteases involved in cytokine cleavage.

*Giardia* has been previously shown to downregulate certain signatures of inflammation both in patients and *in vitro*, which may play a role in the frequency of subclinical infections. High parasite burdens in children have been linked to reduced markers of environmental enteric dysfunction, namely reduced alpha-1-antitrypsin, myeloperoxidase, and neopterin in fecal samples (Dougherty and Bartelt 2022; Giallourou et al. 2023; Rogawski et al. 2018). *In vitro* data shows that cathepsin B cysteine proteases released by the parasite directly degrade CXCL8, a chemokine released by enterocytes during inflammation to promote neutrophil recruitment (Cotton, Bhargava, et al. 2014; Zhu et al. 2021). In animal co-infections with *Clostridium difficile* and colonic biopsies from Crohn’s disease patients, *Giardia* similarly decreases neutrophil activity (Cotton, Motta, et al. 2014).

The potentiation of the IL-10 response in dual stimulation with *Giardia* and LPS may similarly reduce inflammation in the intestine during parasite infection. IL-10 has a significant role in controlling intestinal pathologies; IL-10 knockout mice spontaneously develop colitis and experience significantly worse symptoms when treated with dextran sodium sulfate (DSS) (Jofra et al. 2019; Kühn et al. 1993). We speculate that the parasite is inducing IL-10 as a potential survival strategy to decrease parasite clearance. Up to 40% of children in the Malnutrition and Enteric Disease (MAL-ED) cohort study experienced persistent *Giardia* infections (Rogawski et al. 2017). Interestingly, IL-10 deficient animals do not clear *Giardia* faster than wildtype controls (Dann et al. 2018). However, the *G. muris* used in that study has different infection kinetics than the *G. duodenalis* strain that infects people and was used in our work.

Recognition of pathogens through pattern recognition receptors is thought to be essential in activation of innate immune responses. However, very little is known about the host receptors involved in *Giardia* recognition. We previously reported that Mannose Binding Lectin (MBL) recognized *Giardia* and activated complement (E. Li, Tako, and Singer 2016). In addition, *Giardia* extracellular vesicles upregulate production of pro-inflammatory cytokines by thioglycollate-elicited peritoneal macrophages in a TLR2 and NLRP3 dependent fashion (Zhao, Cao, Wang, Dong, et al. 2021). *Giardia* variant-specific surface proteins similarly signal through TLR4 and TLR2 on reporter HEK293 cells (Serradell et al. 2019). Our new data indicate that MGL1 is aso an important receptor for the development of immune responses in giardiasis. The role of MGL1 has previously been identified in maintaining tolerance to commensal bacteria in the intestine via IL-10 secretion from macrophages (Saba, Denda-Nagai, and Irimura 2009). MGL1^-/-^ animals treated with dextran sodium sulfate (DSS) experience reduced IL-10 expression and increased levels of inflammation in response to bacteria crossing the damaged intestinal layer (Saba, Denda-Nagai, and Irimura 2009). Lamina propria macrophages, in fact, interact directly with glycoproteins on the surface of the commensal *Streptococcus sp*. to release IL-10 (Kurashina et al. 2021).

While we have previously shown that macrophages in the lamina propria proliferate following *Giardia* Infection, the current studies were performed on peritoneal macrophages(Fink et al. 2019). Lamina propria macrophages are more difficult to isolate in the high numbers we needed for stimulation experiments. The two populations likely share some similarities, however, because they occupy the same environment and are rapidly recruited to the intestine during damage (Honda et al. 2021). Additionally, while our studies did not show an upregulation in TNF-□ production due to *Giardia* stimulation alone, others have seen this type of response (Zhao, Cao, Wang, Dong, et al. 2021). It is important to note, however, that these studies used *Giardia* secretory products, as well as thioglycollate-primed macrophages.

While our data indicate that MGL1 is involved in the production of IL-10 by macrophages, the nature of the parasite ligand requires additional study. MGL1 normally binds terminal Galactose, GalNAc, LacdiNAc, and Lewis X and A structures (Vliet, Saeland, and Kooyk 2008). We know that the cyst wall contains GalNAc polymers, but how much if any of this high molecular weight polymer is found in trophozoites is disputed. There appears to be low-level activity of the UDP-GalNAc transferase, the enzyme responsible for GalNAc glycosylation of proteins, in trophozoites, which increases during encystation (Das and Gillin 1996). Others, however, have seen no GalNAc on *Giardia* trophozoite secretory-excretory vesicles, although notably they did find terminal Galactose, another MGL1 ligand (Morelle et al. 2005). Further studies on the MGL1 ligand in *Giardia* are therefore needed.

Asymptomatic *Giardia* infections are very poorly understood. The parasite has been reported to have a fair number of anti-inflammatory effects in the past, which may be contributing to milder symptoms. We and others suggest that the reduction in symptoms may be due to an enhanced IL-10 response from immune cells. It has recently been shown that *Giardia* infection prior to either DSS treatment or infection with *Toxoplasma gondii* leads to a reduction in intestinal inflammation and an increase in IL-10 producing FOXP3^-^ GATA3^+^ CD4 T cells (Sardinha-Silva et al. 2024). In this study, we show that *Giardia* can synergize with bacterial LPS to promote increased IL-10 production from macrophages, a response likely occurring 2prior to the increase in IL-10 from T cells. Increased IL-10 could explain the persistence of *Giarda* in children, as well as the frequent presence of the parasite in control stools (Kotloff et al. 2013; Rogawski et al. 2017).

