## Supplemental Tables for "*Giardia* Increases Macrophage Production of the Anti-Inflammatory Cytokine Interleukin-10 in Response to Lipopolysaccharide via Macrophage Galactose Binding Lectin (MGL1)"

**Supplementary Table 1: Raw TNF-α cytokine production (pg/mL)**

| Cells | Unstimulated | LPS | *Giardia* | LPS + Giardia |
|---|---|---|---|---|
| **WT[i] Pool 1** | 23.7 | 1818.9 | 70.1 | 1821.8 |
| **WT Pool 2** | <15.6[ii] | 1751.9 | 52.5 | 3089.7 |
| **WT Pool 3** | <15.6 | 2402.1 | <15.6 | 3699.2 |
| **WT Pool 4** | <15.6 | 1487.9 | <15.6 | 2452.2 |
| **WT Pool 5** | <15.6 | 3137.1 | 16 | 2870.8 |
| **WT Pool 6** | <15.6 | 4543.8 | 16.4 | 2945 |
| **WT Pool 7** | <15.6 | 1509.6 | <15.6 | 317 |

[i] WT stands for wildtype

[ii] Values below the limit of detection (15.6 pg/mL of TNF-α) are indicated as such.

**Supplementary Table 2: Raw IL-10 cytokine production (pg/mL)**

| Cells | Unstimulated | LPS | Untreated Giardia | LPS + untreated Giardia | Heated Giardia | LPS + heated Giardia | Spun Giardia | LPS + spun Giardia |
|---|---|---|---|---|---|---|---|---|
| **WT[i] Pool 1** | <31.25[ii] | 177.3 | 148.9 | 353.9 | 121.1 | 416.4 | 109.3 | 314 |
| **WT Pool 2** | <31.25 | 44.3 | 50.9 | 287.3 | 75 | 434.9 | 43.5 | 275 |
| **WT Pool 3** | 33.7 | 302.8 | 43.3 | 440.2 | 62.2 | 496.4 | 42 | 418.4 |
| **WT Pool 4** | 47.5 | 322.9 | 54.7 | 384.5 | 91.1 | 517.3 | 68 | 425.4 |
| **WT Pool 5** | 44.1 | 364.8 | 42.2 | 649.3 | 85.8 | 1041.3 | 59.1 | 519 |
| **WT Pool 6** | 66.6 | 346.5 | 38.8 | 627.7 | 67.1 | 1025.4 | 57.7 | 442.9 |
| **WT Pool 7** | <15.625 | 34.1 | 17.3 | 85.3 | 24.5 | 167 | 30.8 | 105.7 |

[i] WT stands for wildtype.

[ii] Values below the limit of detection (15.6-31.2 pg/mL of IL-10) are indicated as such.

**Supplementary Table 3: Raw IL-10 production by macrophages from MGL1$^{+/-}$ animals (pg/mL)**

| Cells | Unstimulated | LPS | *Giardia* | LPS + Giardia |
|---|---|---|---|---|
| **MGL1$^{+/-}$ Mouse 1** | <15.6[i] | 70.7 | 31.6 | 110.2 |
| **MGL1$^{+/-}$ Mouse 2** | 34.9 | 58.4 | 48 | 94.4 |
| **MGL1$^{+/-}$ Mouse 3** | 38.2 | 157.8 | 92.8 | 173.6 |
| **MGL1$^{+/-}$ Mouse 4** | 51.8 | 171.6 | 60 | 138.4 |
| **MGL1$^{+/-}$ Mouse 5** | <15.6 | 96.1 | 13.4 | 223.3 |
| **MGL1$^{+/-}$ Mouse 6** | 18.5 | 291 | 28.5 | 464.1 |
| **MGL1$^{+/-}$ Mouse 7** | <15.6 | 114.6 | 16.4 | 199 |

[i.] Values below the limit of detection (15.6 pg/mL of IL-10) are indicated as such.

**Supplementary Table 4: Raw IL-10 production by macrophages from MGL1$^{-/-}$ animals (pg/mL)**

| Cells | Unstimulated | LPS | *Giardia* | LPS + Giardia |
|---|---|---|---|---|
| **MGL1$^{-/-}$ Mouse 1** | 112.7 | 123.2 | 90.6 | 128 |
| **MGL1$^{-/-}$ Mouse 2** | 94.2 | 366.7 | 103.7 | 405.8 |
| **MGL1$^{-/-}$ Mouse 3** | 97.7 | 467.2 | 79.8 | 309.5 |
| **MGL1$^{-/-}$ Mouse 4** | 126.6 | 381.9 | 111.1 | 334 |
| **MGL1$^{-/-}$ Mouse 5** | <31.2[i] | 220.9 | <31.2 | 261.1 |
| **MGL1$^{-/-}$ Mouse 6** | <31.2 | 341.6 | <31.2 | 345.1 |

[i] Values below the limit of detection (31.2 pg/mL of IL-10) are indicated as such.
